# Automatic Detection and Extraction of Key Resources from Tables in Biomedical Papers

**DOI:** 10.1101/2024.10.15.618379

**Authors:** Ibrahim Burak Ozyurt, Anita Bandrowski

## Abstract

Tables are useful information artifacts that allow easy detection of data “missingness” by humans and have been deployed by several publishers to improve the amount of information present for key resources and reagents such as antibodies, cell lines, and other tools that constitute the inputs to a study. The STAR*Methods tables, specifically, have increased the “findability” of these key resources, but they have not been commonly available outside of the Cell Press journal family. To improve the availability of these tables in the broader biomedical literature, we have attempted to automatically process BioRxiv preprints to create tables from text or to recognize tables already created by authors and structure them for later use by publishers and search systems, to improve “findability” of resources in a larger amount of the scientific literature. The extraction of key resource tables in PDF files by the best in class tools resulted in Grid Table Similarity (GriTS) score of 0.12, so we have created several multimodal pipelines employing machine learning approaches for key resource table page identification, Table Transformer models for table detection and table structure recognition and a new table-specific language model for row over-segmentation to improve the extraction of text in tables created by biomedical authors and published on BioRxiv to around GriTS score of 0.90 enabling the deployment of automated research resource extraction tools onto BioRxiv.

**Author summary:** Tables are useful information artifacts that allow for easy detection of data “missingness” by humans and have been implemented by several publishers to improve the amount of information present for key resources and reagents such as antibodies, cell lines, and other tools that constitute the inputs to a study. To improve the availability of these tables in the broader biomedical literature, we introduced four pipelines for key resource table extraction from biomedical documents in PDF format. Our approach reconstructs key resource tables using image level table detection and structure detection generated table boundary, column (and row) bounding box information together with PDF text alignment. To remedy row over-segmentation resulting from overflowing table cell contents, we introduced a language modeling (LM) based row merging solution where a character-level generative pre-trained transformer (GPT) model was pre-trained on more than 11 million scientific table contents from PubMed Central Open Access Subset (PMC OAS). All introduced pipelines significantly outperformed GROBID baseline while our Table LM based row merging based pipeline, significantly outperformed all other pipelines including our OCR based pipeline.

## Introduction

A table is defined (as of Oxford English dictionary) as an arrangement of numbers, words or items of any kind (content aspect), in a definite and compact form (structure aspect), so as to exhibit some set of facts or relations in a distinct and comprehensive way, for convenience of study, reference, or calculation (function aspect).

Resource tables in scientific papers, such as the STAR*Table [1] denote the use of chemical reagents, antibodies, cell lines, organisms, software tools, instruments, and other experimental inputs to the study. Many of these types of research resources are associated with erroneous reporting practices [2], and have been called out by the community as the main culprits and sources of variability that lead to the reproducibility crisis [3]. The STAR*Table became a very powerful tool in bio-medicine because these simple three column data structures provided the immediate visual stimulus denoting missing information, that might ordinarily be hidden in long paragraphs, encouraging the author to look up identifiers, verify names, and be alerted that an issue may have been reported by their colleagues about the resource which may impact their conclusions. Before these tables were implemented, the percentage of antibodies that was identifiable hovered between 10 and 30% precluding direct replication of 70 to 90% of antibody-using papers, but after the implementation of the STAR*Methods table in the journal Cell, over 95% of antibodies were identifiable [4]. The ability to determine which reagent is used in a particular study is a major source of irreproducibility in the scientific literature [5], and a source that should be “easy to fix” as most authors simply need a quick reminder to pull the appropriate information from laboratory records into the paper [6]. Therefore these tables by their simplicity are highly effective tools for authors to improve transparency of scientific work. Some characteristics and issues of key resource tables found in scientific papers are summarized in Fig 1.

**Fig 1.**
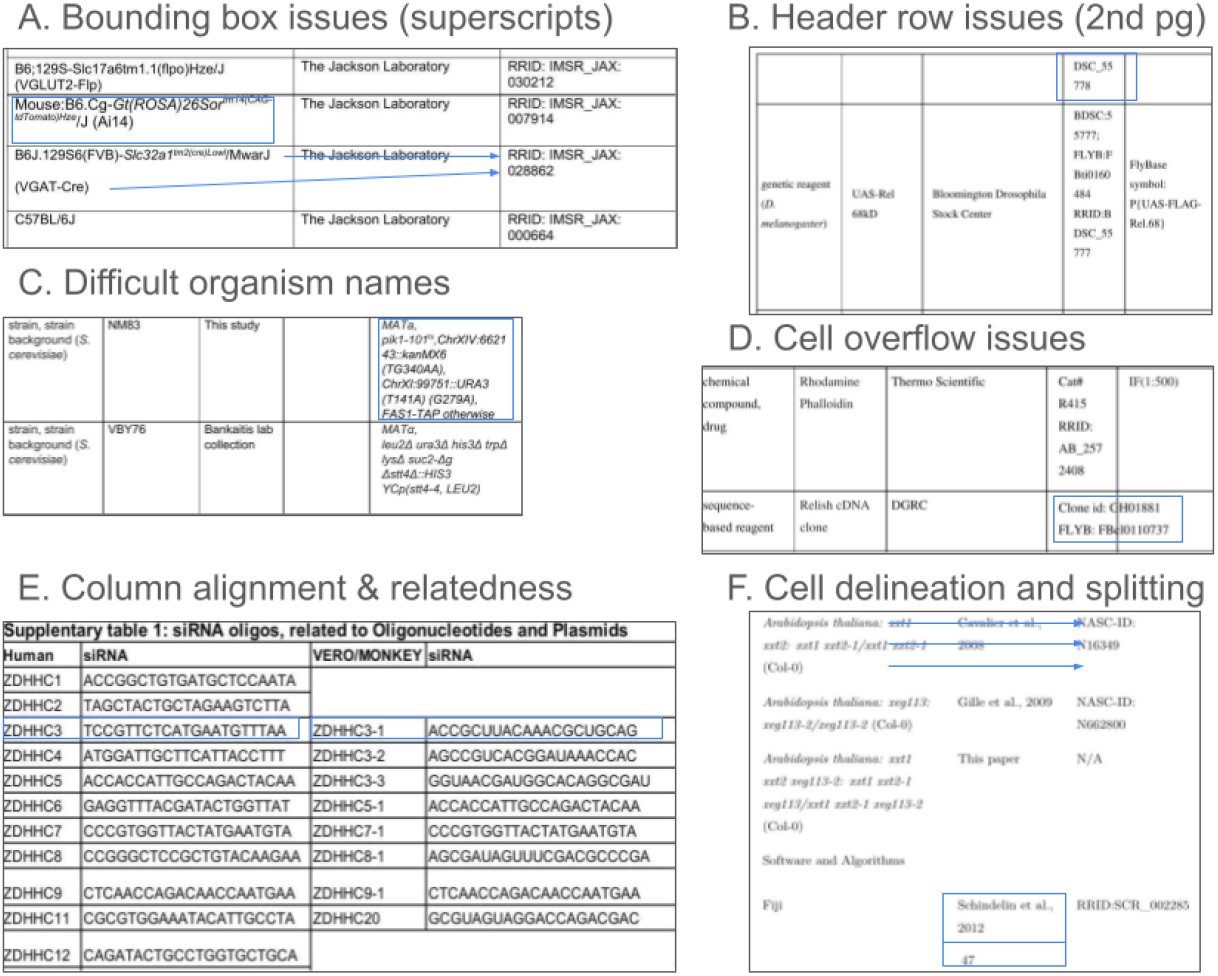
Common issues with key resource tables.

The ability of STAR*Tables to reduce omitted resource information in the scientific literature is directly related to their use. Unfortunately, most journals do not enforce a standard resource table due to the lack of manpower to police this type of policy (personal communication). Preprints submitted to BioRxiv, the major preprint server for biology, are not checked by editorial staff except to determine if the manuscript is obviously not a scholarly work and thus may be the place where this information omission is most acute, but as preprints are not yet published and may be commented on, this is a perfect place for intervention that might increase the number of these resource tables in the bio-medical literature. To help authors of BioRxiv preprints improve their manuscript by quickly reminding them that some information about some resources may be missing, we undertook the task of automatically creating resource tables for BioRxiv preprints. With detection and text processing we should be able to provide the information to authors as automatically created tables and encourage them to look over the result and fix any errors of omission that were introduced by either the process of writing the manuscript or the extraction of text into tables. However, for authors who already include a resource table in their preprints, we should be able to detect them and display the results. The BioRxiv team creates images of tables reducing their ability to be operated on because the task of coding tables into a more interpretable format is too resource intensive (Richard Sever, personal communication). This task requires that tables created by authors in arbitrary formats need to be found and correctly interpreted as to the table structure and cell values.

Wang [7] provided a formal model for the logical structure of a table. A table consists of entities, the basic data that it displays, and labels, the auxiliary data used to locate the entities. The labels are further classified into categories that form a tree structure. Based on this model, a table is a mapping of entries with frontier label sequences from a tree-structured set of categories. During table construction, the logical structure of a table is converted into a two-dimensional grid structure for representation. Table data information extraction is the reconstruction of the underlying implicit logical model of a table from its representation. Viewed from the perspective of natural language processing (NLP), this task is made more demanding by two factors. First, there are long distance relationships between table labels and entities that do not fit human language syntax and semantics. Second, there is almost no redundancy in the entities and labels to learn syntax and semantics. Attacking the problem from the representation side using computer vision techniques is another approach to infer the logical structure from the layout. However, this approach requires optical character recognition (OCR) to convert the content of detected table cells back to text, resulting in character level errors (even an OCR method with 99% accuracy will have a wrong character every 100 characters on average). To illustrate why even this low an error rate may be problematic we can imagine that we are examining a PDF file of an invoice which must be paid, but one of the numbers is incorrect without any indication as to which one is incorrect leading to potential problems in the case where the error is in something like the hundreds place. Nearly any character can be found in a catalog number or an organism name, and even one mistake in 100 will invariably lead to a different item.

Figure 1 summarizes some of the most commonly encountered issues in resource tables, and they include problems such as places where superscripts or other characters are very close to bounding boxes (A), creating potential errors in OCR parsing because the bounding box may be interpreted as part of the character. Some tables that are present on more than a single page may only have one header row and some cells may be broken between page one and page two causing errors in data stitching, in the case shown in (B) the first row on page 2 of the table contains a part of the identifier that is associated with the fly on the previous page. Some organism names are fairly long and may have multiple lines with characters that are more similar to math symbols than English names (see C). In a relatively common case for BioRxiv preprints authors sometimes do not notice or fix cases where the contents of a single cell overruns that cell, and in at least one case the overflow is not visible because it is covered by white box in the next cell, and only shows up in the text extracted version of the table. Tables are also not all structured in the same way, and we cannot assume that a row always means that the information content is about the same entity. Indeed in part (E) the table contains a set of oligonucleotides that are similar between humans and monkeys, but they are not the same reagents. Many tables do not contain any bounding box information, which makes it more difficult to know where one cell begins and another ends, in part (F) the bounding box at the bottom contains a reference and a page number which might be stitched together, and information in two cells that take three rows in the left most cell, two lines in the middle cell and two lines in the right most cell. Knowing that the information should be stitched together not one row at a time but three rows, two rows and two rows is not trivial for an OCR or for text extraction systems.

## Related Work

While early approaches for table extraction where mostly rule and heuristics based [8], [9], current approaches [10], [11], [12] rely on deep learning based object detection on images of document pages. These approaches typically use convolutional neural networks (CNN) and/or recurrent neural networks (RNN) requiring a large amount of labeled training data.

Preparation of the training data for image based approaches involve labeling of bounding boxes for each table cell for a large number of tables which is a very resource intensive task. PubMed Central provides millions of articles and preprints in NISO Journal Archiving and Interchange (JATS) XML format also encoding tables occurring in the corresponding papers. Besides JATS XML files corresponding PDF are also provided. By associating the tables encoded in JATS XML files with their corresponding PDF character locations, a large set of labeled table data can be generated. Using this approach, training/evaluation sets are generated [12], [13] for table extraction.

In scientific and science domain GROBID [14] is currently the most popular tool for extracting, parsing and re-structuring PDFs into structured XML/Text Encoding Initiative (TEI) encoded documents including tables. Recently, Table Transformer [13] models trained on tables extracted from the PMC OAS are introduced.

## Materials and methods

### Systems Overview

Towards the goal of automatic detection of key resources from tables in PDF preprints and supplementary documents, four pipelines are introduced as summarized in Fig 2. The key resource candidate detection module is common in all the pipelines and selects the candidate pages in a paper to be used in the following pipeline module which uses Table Transformer table detection (TD) model to detect table bounding boxes, followed by cell bounding box prediction via the Table Transformer table structure detection (TSR) model. Pipelines A, B and D are hybrid multimodal approaches relying image level models to detect table, column and (for model B) row boundary information followed by extraction of corresponding cell text from PDF. Extraction of the text from the PDF for each cell identified by the TSR bounding box information (whether it is just column range or both column and row range information) is done by aligning the character bounding boxes and applying canonicalization steps on the candidate strings which are detailed in following sections. Column ranges derived from TSR model predicted cell boundary boxes were observed to be less error prone than the row ranges since separation between table columns are almost always more than the separation between table rows in table images. Based on this observation, pipeline A only uses column ranges derived from TSR model cell boundary box predictions resulting in over-segmentation of the table rows with overflowing cells. Pipeline B uses both column and row ranges derived similarly from TSR model predictions. To remedy the row over-segmentation problem resulting of spurious rows for overflowing cells, in pipeline D a language model based approach is introduced for pipeline A to merge over-segmented table rows reliably. Pipeline C works on only image modality where the text from the TSR predicted bounding box images were extracted using optical character recognition.

**Fig 2.**
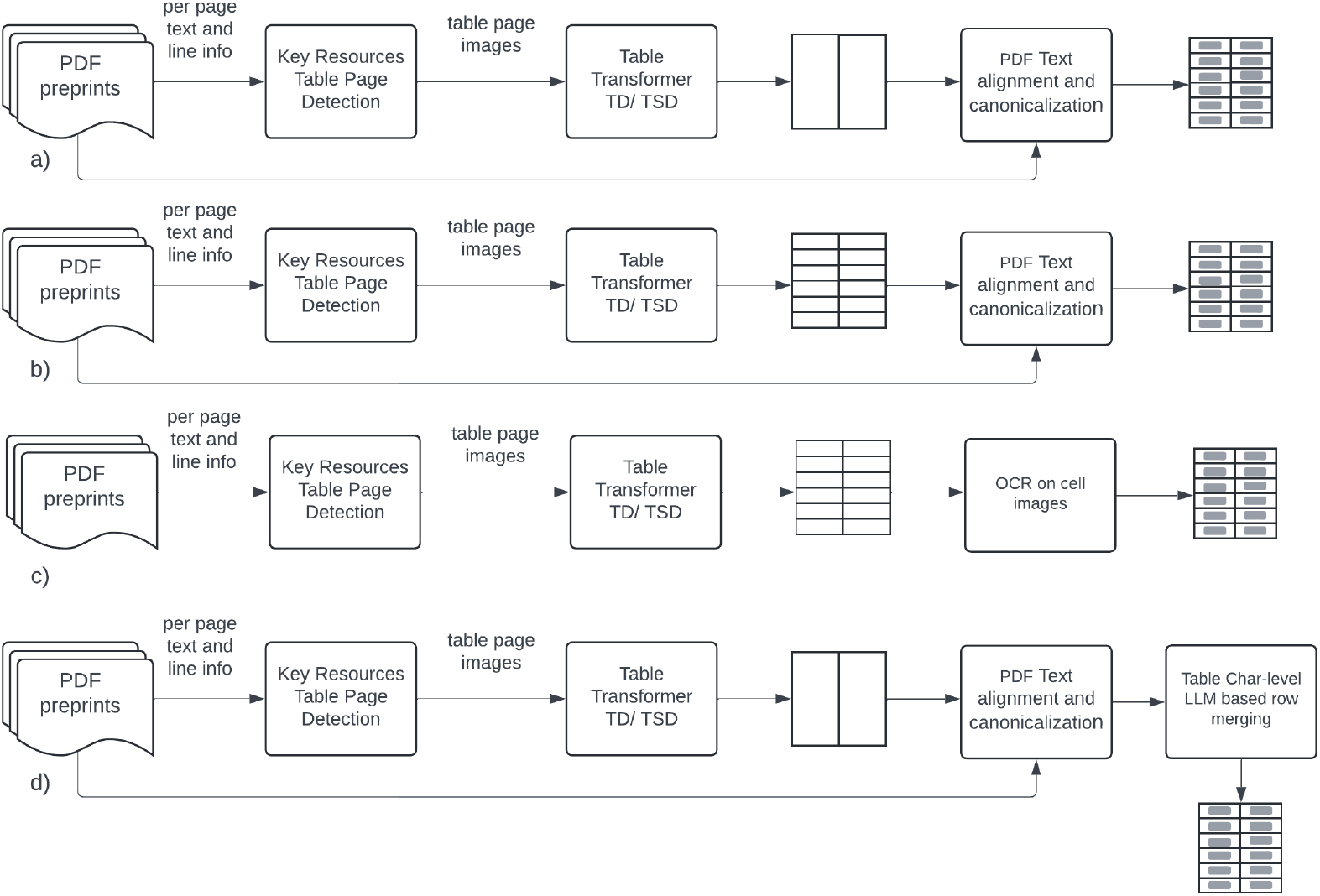
Key resource table reconstruction pipelines. a) Pipeline A (using only TSR column location info) b) Pipeline B (using both TSR column and row location info) c) Pipeline C (fully image based using OCR for cell contents) d) Pipeline D (using only TSR column information together with character level table language model based row merging)

### Detection of Resource Table containing Pages in PDFs

Since we are interested in detection of a specific type of table, namely key resource containing tables, PDF pages that contain key resource tables need to be detected as the preliminary step before the detection of tables, their structures and finally reconstruction of their content. In PDF, each character has its own bounding box [*x*_*min*_, *y*_*min*_, *x*_*max*_, *y*_*max*_]. Also, shapes such as grid lines of a table are represented as polygons with vertex coordinates specified. Since not all tables have a grid and other document structures such as figures are also constructed from primitive polygons, by themselves grids are not indicative of a table occurring in a table when they are detected. Identifying the pages that potentially contain key resource tables is approached using a stacked generalizer [15] ensemble classifier where the predictions of a classifier is used as additional features for a second classifier making the final decision.

The first level classifier operates on local line level features to predict whether the current line of a PDF document page is inside a key resource table or not. For this classifier, a long short-term memory (LSTM) [16] based neural network is employed using both text based and structural features. Using a windowing approach, the tokens of the current line and any previous line constitute the local textual context. For structural features, any vertical and horizontal lines as encoded in PDF primitive graphics operations for the current and any previous line are used as indicator features. The current page number is also used as a feature. As token embeddings, GloVE [17] word vectors trained in house on PubMed abstracts were used. In the neural network, the textual context embedded as pre-computed GloVE word embeddings is encoded using LSTMs and concatenated via the other structural indicator features for a final logistic regression layer. To compensate for the imbalance in the number of lines of text in a table versus not, the errors in positive class are penalized 50 times more than the negative class errors during training.

The second level classifier is a linear support vector machine (SVM) operating on the text content of the whole page together with the number of in table lines predicted by the first level classifier. By using a very common text classification assumption, bag of words (BoW) assumption, the unique words in the training corpus is represented by their TF-IDF score to model their relevance in the current page. Since the number of pages that are not key resource tables is at least an order magnitude more than key resource containing tables, to compensate for the imbalance in negative and positive classes, errors in the positive class are penalized five times more than the negative class in the objective function for the SVM classifier.

To train and test the introduced system, 57 BioRxiv preprints containing key resource table(s) were manually labeled by extracting their text content including page numbers and marking the beginning and ends of each resource table or pages for multi-page spanning tables. The generated labeled preprints were randomly split to a training set of 47 preprints and 10 preprints for testing.

### Table Detection and Table Structure Detection using Table Transformers

Table Transformer [13] models were trained on 948K tables from PMC OAS corpus (RRID:SCR_004166). JATS XML tables cell contents are aligned with the corresponding PDF text boxes via Needleman-Wunsch algorithm [18]. A canonicalization algorithm to correct over-segmentation errors in the structure annotation of a table (merging adjacent cells under certain conditions) was also introduced [13]. A DETR [19], a CNN-Transformer object detection model, with a ResNet-18 backbone and few frozen first layers trained on Imagenet data was fine-tuned for table detection (TD) and Table Structure Detection (TSR). For this work, we had used Table Transformer models available from Hugging Face (RRID:SCR_020958), https://huggingface.co/docs/transformers/main/en/model_doc/table-transformer).

### Object Detection to PDF Alignment and Canonicalization

To facilitate the alignment of the Table Transformer TD and TSR bounding boxes with the corresponding PDF document text bounding boxes, first the bounding box coordinates are scaled to PDF page coordinates as the Table Transformer model pre-processing involves image scaling. This is followed by column and row range estimation and heuristic row merging steps described in the following sections.

#### Column and Row Range Estimation

The effective number of columns of a table is predicted as the median of the number of row cells in the TSR bounding boxes. After that for each column, the lower bound is the minimum *x*_*min*_ value and the upper bound is the maximum *x*_*max*_ value of all the detected cell boundary boxes for that column plus about 1 pixel additional space to minimize column boundary alignment errors. The bound of the last column from the Table Transformer TSR bounds is expanded by two average character widths to compensate for occasional clipping of last characters due to tight bounding box errors.

For pipelines without row information from Table Transformer TSR model, row ranges are are estimated by first grouping all character position bounding boxes in the PDF document by their top *y* coordinate followed up by merging all characters that are connected horizontally resulting in text lines. Since for superscript and subscript character bounding boxes have different top *y* values than their baseline, this operation creates spurious text lines which are handled by a separate canonicalization step.

#### Canonicalization (Row Merge Phase)

Using only the more robust table column range information estimated based on the Table Transformer TSR model results in over-segmentation of the table rows. Besides that, any superscripts/subscripts are raised/lowered from the baseline text in PDF extracted text lines. To decrease the number of spurious rows generated, following heuristic rules are used;

1. if a row text range overlaps 70% or more with the closest row rectangle range, merge both rows (mostly for subscripts and superscripts)
2. If a row has empty cells and a row above without empty cells and row text range overlaps 50% or more with the closest row text range, merge both rows

While these rules work in a lot of times, they cannot handle all cases.

### Optical Character Recognition for Table Cells

For the Optical Character Recognition, OCR, based pipeline C, the table cell content images were extracted based on the predicted bounding boxes from the TSR model. For OCR, we have used the open source Tesseract [20]. Each cell image was first converted to gray scale and contrast enhanced before passing it to the Tesseract OCR model. We used the ‘Assume a single uniform block of text’ option of Tesseract to minimize OCR errors.

### Learning the Language of Scientific Tables

Human language generation is traditionally modeled as a joint probability distribution of a sequence of conditional probabilities of tokens termed language model [21]. Based on this model, given an sequence of symbols (*s*_1_, *s*_2_, …, *s*_*n*_) such as words or characters in a sentence, the conditional probability of generating the next word *s*_*k*_ depends on the previous context expressed as *p*(*s*_*k*_|*s*_1_, *s*_2_, …, *s*_*k*−1_). Current advances in deep learning architectures, specifically Transformer architecture [22] allowed increasingly more expressive models of these conditional probabilities by increasing the width and the height (number of transformer layers) of the underlying neural network.

For training corpus preparation, the PMC OAS June 2024 set of full papers in JATS XML format is used. From these papers, 11,467,759 tables are extracted by parsing their corresponding JATS XML file. From each table, the content of each cell including the header rows are used to build our pre-training corpus of about 1.7 billion tokens.

Since a table cell usually contains condensed information articulated with a few words or numbers and with the ultimate goal of being able to predict if the content in the potentially over-segmented next row under the current cell is the continuation of the current cell, a language model at character level instead of the most common word piece level is used. Thus, an auto-regressive generative language model is trained to predict the next character given the characters up to the next character for all the cell contents of all tables available in PMC OAS papers.

As the language model, a generative pre-trained transformer (GPT) [23] architecture with decoder transformer layers using causal multi-head self attention is selected. Causal self attention, unlike the transformer encoder models such as BERT [24], only attends to the tokens before the predicted next token. A six transformer layer GPT model with 6 attention heads and 384 dimensional embedding vectors is used to model the language of the table contents of the biomedical papers. The resulting model has 15.6 million parameters and can handle sequences up to 256 characters. A vocabulary of 11,946 unique characters and ‘<EOS>‘ special token to indicate the end of sequence (i.e. end of table cell content) is used. The language model is pre-trained on a RTX 4090 24GB GPU with a batch size of 256 for 250,000 steps in 8 hours.

### Learning to merge overflowing table cell content

A solution to the table row over-segmentation problem for the extracted table data reconstruction is learning to classify if the contents of two vertical neighboring cells should be merged or not. Being able to do this requires domain knowledge such as recognizing different representations of organisms, cell lines, antibodies or genetic sequences, plasmids and ability to exploit syntactic and semantic level language clues such as a hyperlink spread across multiple rows. Our hypothesis is that a character level language model pre-trained on the contents of a large number of table cells will implicitly learn the syntactic and semantic properties of the language used in biomedical domain tables to represent results and data characteristics. Given a table of *C* columns and *R* rows, where a cell content of the ith row and jth column is denoted as *c*_*ij*_, the binary classification problem for a single table can be stated as *f* : **X** → *y* = *{*0, 1*}* where **X** ∈ *{c*_*ij*_| *<EOS >* |*c*_*i*+1,*j*_, 0 *<*= *i < R* − 1, 0 *<*= *j < C}*. Here *y* are binary labels indicating whether *c*_*ij*_ and *c*_*i*+1,*j*_ should be merged or not and | is the concatenation operator. For transfer learning, the head layer of the character level table LM is replaced with a single sigmoid neuron and the rest of the model is initialized from the weights of the character level table LM.

### Generating simulated table cell overflow dataset

In order to train the supervised cell merge classification, a large labeled training set of key resource containing tables is necessary which is costly and time consuming to annotate. However, cell merge requiring scenarios can be simulated from the contents of JATS XML represented tables in PMC OAS. In order to do this, first the key resource containing tables from more than 11 million tables in PMC OAS dataset needs to be selected in a reliable way. Since RRIDs are used increasingly with key resource tables, we filtered PMC OAS tables for ‘RRID:’ prefix resulting in 13,664 tables to create a simulated cell merge training set.

To simulate one or more cells overflowing into multiple rows, all tables having at least one row over the size 90 characters (maximum number of characters to fit a single line on a page with reasonable font size) are selected. For tables with less than 100 max character total width as defined as the length of all its column contents of its longest row, a 80 character width maximum is used. The allowed total row width is divided to each column in proportion to their average column width plus the standard deviation of the column’s width. This process allocates additional space for columns with larger variation of widths to simulate the way the authors select column widths to minimize column spillovers. After the column widths are determined, any cells that do not fit their allocated column width will be candidates for positively labeled cell merge pairs. Negative training examples can be generated from non overflowing cells in neighboring rows of the same column. A simulated overflowing cell is split at space characters if possible to as many rows necessary to represent the content.

### Row Merge Prediction

The 13,664 key resources table corpus is randomly divided in a 90%/10% training / testing set. Using this approach, 2.8 million simulated training instances and 311,998 testing instances are generated. The binary classification model is trained for two epochs with Adam optimizer using a learning rate of 2e-4. The model achieved an accuracy of 98.7% on the test set of key resource tables.

Since the introduced key table resource cell merger classifier only predicts whether two rows in consecutive cells of the same column should be merged, a maximum voting based approach is used to decide which neighboring table rows to merge. To achieve this for any pair of table rows, the cell merger classifier was applied to each column cell pair and a row score was generated by summing the predicted column cell merge probabilities and finding the average value. If the row score is over 0.5 then the neighboring rows were merged.

### Key Resource Table Extraction Gold Standard Set Construction

Evaluating of the introduced approaches requires a gold standard set of reconstructed tables to test against. Due to the lack of such a resource, April 2024 collection of 4652 BioRxiv preprints were downloaded. Preprints containing the keywords ‘RRID’ and ‘antibod’, common keywords potentially indicative of key resources, were selected resulting in a set of 1655 candidate preprints which are processed by the table extraction pipeline A resulting in 302 preprints with key resource tables detected. Out of this set 50 preprints were randomly selected for manual correction of over-segmentation errors to generate the gold standard set. The final gold standard set contains 100 tables in 46 BioRxiv preprints and was used in evaluation of the key resource table extraction pipelines introduced.

### GROBID Baseline Table Extraction

As a baseline, GROBID [14] system is also tested against our constructed gold standard table set. The corresponding PDF documents of the gold standard tables were processed via locally installed latest (version 0.8.0) GROBID server followed by the parsing of the recognized tables out of the generated XML/TEI documents. After that, the GROBID recognized tables were aligned with the gold standard set by vocabulary overlap of the GROBID table with the corresponding gold standard table. A vocabulary overlap over 40% is considered as a threshold for table alignment to maximize the inclusion of partially extracted tables by GROBID.

### Evaluation Metrics

To evaluate both the topology and content of tables extracted from biomedical paper/supplementary document PDFs, Grid Table Similarity (GriTS) [25] measure is used. GriTS represents the ground truth and predicted tables as matrices and computes two dimensional most similar substructures among these matrices as shown in Eq 1. taking fully into account the two dimensional structure of tables and addressing the cell topology, location and content in a unified manner. Here Ã and 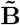 denote the substructures, a selection of rows and columns aligned between two table matrices **A** (ground truth) and **B** (predicted). Three different similarity functions *f* are defined for cell topology, content and layout similarity between the ground truth and predicted table similarity. The cell content and topology similarities are the most relevant for the key resource table extraction.

GriTS_Cont_ measures the similarity of the layout and content of cells, while GriTS_Top_ measures the similarity of the row and columns that each cell occupies in the grid between the ground truth and predicted table. GriTS measure can be interpreted like the commonly used *F* measure which is the geometric mean of recall and precision. Precision and recall for GriTS are defined in Eq 2 and Eq 3, respectively. The similarity of cell contents between the the aligned ground truth cell and predicted cell is calculated as the ratio of longest common string sequences to the length of the ground truth cell content string. The similarity of the aligned ground truth and predicted table substructures is estimated by the intersection-over-union (IOU) of the aligned bounding boxes.

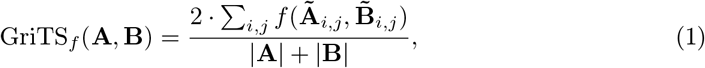

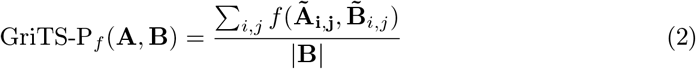

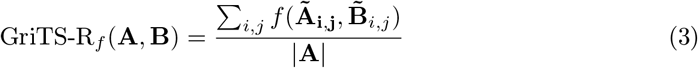

## Results and Discussion

### Table Detection Performance of the Table Transformer on BioRxiv Preprints

To evaluate the performance of the table detection via Table Transformer TD model, 1000 randomly selected preprints (between 2019 and 2023) are further filtered to a set of 143 PDFs (paper body and supplementary files) containing the RRID prefix. After converting to an image, Table Transformer TD model is applied to each page of the 143 PDF documents resulting in 2626 table candidates for which the bounding box content is saved as a separate image for curation. A single annotator looked through each table candidate image and decided if the bounding box contents is an actual table or not. Out of 2626 table predictions only 356 were actual tables with an accuracy of 13.6%. Majority of the errors seem to fall into one of these categories 1) pages with line numbers (a common occurrence in preprints) 2) numbered references 3) figure captions 4) first page author lists 5) other kinds of lists. However, if a page contains any table, detection rate is much higher as reported in [13]. This result demonstrates the need for detection of key resource table containing pages before applying Table Transformer TD model.

### Key Resource Table Page Candidate Detection Performance

The test performance of the key resource table candidate page detection systems are summarized in Table 1. Single BoW features only SVM achieves very high precision but lower recall rate. The two level stacked generalizer model performs slightly better by increasing recall at the expense of some precision. To evaluate model in a larger setting, 200 new randomly selected preprints were passed through the stacked generalizer model and results were curated by student curators. The precision showed a slight drop while the recall remained the same.

**Table 1.** Test performance of key resource table page candidate detection systems.

| Model | Precision | Recall | $F_1$ |
| --- | --- | --- | --- |
| BoW only SVM | 100% | 67% | 80% |
| Stacked Generalizer model | 92% | 73% | 81% |
| Stacked Generalizer model over larger test set | 84% | 67% | 75% |

### Key Resource Table Extraction Performance

The table content and topology extraction performance of the four table extraction pipelines introduced together with the GROBID baseline against the 100 gold standard key resource tables is summarized in Table 2. All of the introduced pipelines outperform GROBID baseline with a significant margin. GROBID was only able to detect 19 of the 100 gold standard tables resulting in low GriTS table similarity scores. The best performing pipeline, pipeline D with Table LM based row merging significantly outperforms the OCR pipeline (pipeline C) using two-tailed t-test with a p-value 0.01. Out of the introduced pipelines, the pipeline that uses both column and row bounds information from the Table Transformer TSR model was the worst performing model. This is mainly due the distances between the rows being usually much smaller than the column distances, resulting in row boundary errors and cropped bounding boxes leading to PDF text to table cell alignment issues. Due to the row over-segmentation, the column information only pipeline A showed lower performance than the OCR based pipeline C, especially for content and topology precision namely GriTS-P_*cont*_ and GriTS-P_*top*_. Pipeline D with character level Table LM for row merging remedied the row over-segmentation issue of the pipeline A relying on learned syntactic and semantic properties of the language of scientific tables.

**Table 2.** Test performance of key resource content extraction pipelines.

| Pipeline | GriTS<sub>cont</sub> | GriTS-P<sub>cont</sub> | GriTS-R<sub>cont</sub> | GriTS<sub>top</sub> | GriTS-P<sub>top</sub> | GriTS-R<sub>top</sub> |
| --- | --- | --- | --- | --- | --- | --- |
| Grobid | 0.1162 | 0.1190 | 0.1380 | 0.1341 | 0.1381 | 0.1643 |
| A (TSR Col) | 0.7450 | 0.6804 | 0.8907 | 0.7842 | 0.7103 | 0.9565 |
| B (TSR Row) | 0.5893 | 0.7879 | 0.6196 | 0.6753 | 0.9091 | 0.6196 |
| C (Full OCR) | 0.7553 | 0.7483 | 0.7798 | 0.8496 | 0.8419 | 0.8805 |
| D (TSR Lang) | **0.8981** | **0.8898** | **0.9172** | **0.9262** | **0.9158** | **0.9504** |

GriTS<sub>f</sub>, GriTS-P<sub>f</sub> and GriTS-R<sub>f</sub> are defined in Eq 1, Eq 2 and Eq 3, respectively. Pipelines include Grobid only, Pipeline A (using only TSR column location information), Pipeline B (using both TSR column and row location information), Pipeline C (fully image based using OCR for cell contents), Pipeline D (using only TSR column information together with character level table language model based row merging)

While OCR based pipeline C was the second best performing pipeline, OCR usually results in point errors within recognized words which could make them not recognizable. This is especially a problem for key resources which are required to be identifiable for the reproducibility of a study. For example, the following nucleotide sequence ‘GCACTTCATCCTTTGG G ‘ recognized by the OCR misses a portion of the full sequence ‘GCACTTCATCCTTTGGTTTTG’. OCR can mangle the key resource name, e.g. ‘Alexa antl-Maddil IQ’ instead of ‘Alexa anti-Rabbit IgG’. OCR erros are most detrimental if they occur in a key resource identifier such as a catalog number, e.g. ‘alt 1/404/6’ instead of ‘ Cat# 1745478’. However, the pipeline D with Table LM based row merging extracts the cell contents from PDF text contents always resulting in correct text in the detected table cells.

## Conclusion

While tables are a key visual representation of text and numerical data that are highly useful for human beings because they provide immediate feedback to authors about missing information; however, tables are not easy to extract by conventional approaches. The fact that tables generally contain non-duplicate information in a format that is not English, per se, meant that optical character recognition approaches were insufficient because any error of a character could not be corrected based on context of the character. If, for example, a “g” becomes an “a” in an English word, the word can be directed as being misspelled, but no such misspelling detection is possible when the same error is made in a catalog number or in an organism name where symbols superscripts and subscripts are very common, for example “Pry[+t7.2]=hsFLP12, y[1] w[*]; Pw[+mW.hs]=GawBap[md544] Pw[+mW.hs]=GawBptc[559.1]/CyO, Pw[+mC]=ActGFPJMR1; Pw[+mC]=UAS-TagBFP9D”. Please note, there is little syntax that can be used and it is uncertain for everyone who is not a curator that this is a fly name.

We introduced four pipelines for key resource table extraction from biomedical documents in PDF format. Our approach reconstructs key resource tables using image level table detection and structure detection generated table boundary, column (and row) bounding box information together with PDF text alignment. To remedy row over-segmentation resulting from overflowing table cell contents, we introduced a language modeling (LM) based row merging solution where a character level GPT model is pre-trained on more than 11 million scientific table contents from PMC OAS. All introduced pipelines significantly outperformed GROBID baseline while our Table LM based row merging based pipeline, significantly outperformed all other pipelines including our OCR based pipeline.

### Limitations of this Study

We acknowledge that several key limitations of our current study are present. Due to time consuming nature of the annotation task, the gold set is also relatively small thus may not be representative of all types of resource tables. Additionally, the table extraction code may not generalize for other types of tables, such as statistical tables, because we have trained our tool to specifically detect resource tables. To adapt the pipelines to different type of tables, the table page detection classifier needs to retrained with annotated pages containing relevant tables. Also, the row merger classifier needs to be fine-tuned with synthetic training data generated from relevant type of table contents.

## Supporting information

The key resource extraction pipeline code together with all the annotated data (including gold set used for evaluations) can be found at https://github.com/SciCrunch/key_resource_table_extractor. The Table LM pre-training and row merger fine-tuned classifier code is available at https://github.com/SciCrunch/table_lm. The Table LM model and row merger classifier model can be found at Zenodo (https://doi.org/10.5281/zenodo.13924310). The GLOVE word/phrase embeddings used for key resource page detection classifier are available from https://doi.org/10.5281/zenodo.13924223.

## Acknowledgments

We would like to thank Amit Namburi and Alexander Parker for their contribution to the curation of elements and tables, respectively. We would like to thank Mr. Peter Eckmann for his help in extracting data from BioRxiv. The authors would like to thank the National Institutes of Health [GM144308]. This project has also been made possible in part by grant 2022-250218 from the Chan Zuckerberg Initiative DAF, an advised fund of the Silicon Valley Community Foundation. The funders had no role in study design, data collection and analysis, decision to publish, or preparation of the manuscript.

